# Quantifying Color Vision Changes Associated with Cataracts Using Cone Contrast Thresholds

**DOI:** 10.1101/2020.06.27.175570

**Authors:** Urmi Mehta, Anna Diep, Kevin Nguyen, Bryan Le, Clara Yuh, Caroline Frambach, John Doan, Ang Wei, Anton M. Palma, Marjan Farid, Sumit Garg, Sanjay Kedhar, Matthew Wade, Kailey A. Marshall, Kimberly A. Jameson, M. Cristina Kenney, Andrew W. Browne

**Author notes:** **Corresponding author:** Urmi Mehta.

## Abstract

**Purpose:** The cone contrast threshold (CCT) test quantified color vision changes in subjects of all ages and those undergoing cataract surgery.

**Methods:** Twenty-four healthy volunteers from two cohort studies performed CCT using best corrected visual acuity, filters, mydriasis, and pinhole correction. Retrospective cross-sectional study of patients seen in eye clinics evaluated the relationship between age and color vision, and age and lens status in 355 eyes. Lastly, 25 subjects performed CCT before and after cataract surgery.

**Results:** CCT scores were most reliable in the non-mydriatic condition without pinhole correction. Progressively dense brown filters produced small but significant reductions in S-cone sensitivity. Linear regression analysis of phakic subjects showed a decline for all cone classes with age. Rate of decline was greater for S-cones (slope (95% CI) = −1.09 (−1.23, 0.94)) than M-cones (slope (95% CI) = −0.80 (−0.95, −0.66)) and L-cones (slope (95% CI) = −0.66 (−0.81, - 0.52)). CCT scores increased for S-cones but reduced for L- and M-cones in pseudophakic subjects compared to phakic patients. CCT scores after cataract surgery increased for S-cones, M-cones, and L-cones by 33.0 (p<0.001), 24.9 (p=0.001), and 22.0 (p=0.008).

**Conclusions:** CCT assessment allows for clinically practical quantitation of color and contrast vision improvement after cataract surgery and aging patients who note poor vision despite good visual acuity.

**Translational Relevance:** CCT testing, historically used in research, is now a clinically practical tool to evaluate age and cataract related changes in color and contrast vision and routinely quantify vision beyond black and white visual acuity testing.

## Introduction

Color vision is an integral part of visual function that gradually decreases with age^1-3^. However, visual fields and acuity are the only elements routinely measured or considered in clinical trials. Senescent changes, including senile miosis and increased density and opacity of optical media^2,5^ diminish the amount of light reaching the retina where visual transduction initiates visual acuity, contrast sensitivity, and color vision ^6^. The most common etiology for reduced media clarity is lenticular opacification. With age, the translucent lens begins to yellow and harden toward brunescence, increasing its absorption of short-wavelength visible light, causing acquired tritan deficits ^7-9^. Increasing lens density and the eventual progression to cataracts increase forward light scattering, reducing retinal illumination, contrast sensitivity, and visual acuity while increasing glare^2^.

Most studies of age-related changes in color vision were performed using investigational color vision assays that are not routinely available in clinics. The anomaloscope, considered the quantitative “gold standard” for assessing color vision deficiency (CVD), is a matching test that quantifies abnormalities along the deutan and protan (red-green) axis^14^. Extensive training to administer, high equipment cost, and long test duration yield anomaloscopy a rarely used tool outside research settings. Instead, Ishihara and Hardy-Rand-Rittler (HRR) pseudoisochromatic plates (PIPs), which ask subjects to identify colored numbers or shapes against multi-colored backgrounds on a number of plates with increasing difficulty for object recognition, are employed because they are cost effective and easy to administer. Although all PIPs provide quantitatively discrete data, they provide little information about the severity or extent of CVD^16^.

The Farnsworth-Munsell 100-Hue (FM-100) arrangement test is more discriminative than PIPs. It requires subjects to arrange 85 different colored cards in 4 different groups by order of their hue^17^. Recent adaptations to this test into software have improved previous setbacks, such as length of assessment time and wear and tear to physical pigmented cards leading to hue confusion^18^. However, FM-100 correlates significantly with nonverbal intelligence ^19^ and subjectively determines the degree of CVD^14^. Furthermore, performance complexity and long test duration yield it impractical in routine clinical practice.

Computerized programs, like the custom system used by Fristrom and Lundh, which displayed colored letters on a computer monitor, found a quantifiable negative impact on color sensitivity in subjects undergoing cataract surgery that was greatest for the tritan axis^2^. These prior studies using custom computerized testing or traditional color vision tests are either not broadly available, clinically practical, or provide sufficient and reliable information.

The more recent introduction of the cone contrast threshold (CCT) test answered the challenges posed by traditional color vision tests by allowing for faster and easier testing while producing accurate quantitation and characterization of CVD ^16^. The original cone contrast test by Rabin (RCCT) presented a randomized series of colored letters visible to a single cone type in five steps of decreasing contrast to determine an interpolated contrast threshold ^16,20^. Using RCCT, Fujikawa et al found a gradual decline in CCT scores with age in phakic subjects that increased back to normal values in pseudophakic subjects^10^.

CCT testing differentiates itself from prior testing modalities because it rapidly yields continuous quantitative data for all 3 cone classes. Recent advances in precise computer-based testing and system affordability have yielded logistically practical and readily available clinical visual function test devices to quantify and report quantitative changes in color and contrast vision. This work presents findings from a systematic investigation of media opacity on color vision using a conventional and clinically practical CCT system, ColorDx^®^ CCT HD^®^ (Konan Medical, Irvine CA). CCT HD was a collaborative development under the Cooperative Research and Development Agreement with the US Air Force, School of Aerospace Medicine, Operational Based Vision Assessment Team at Wright Patterson Air Force Base to provide increased reliability and resolution of color vision testing with an expanded low-contrast range not available with the original RCCT. Precision pilot assessment was the original focus of CCT HD. However, efficient and highly granular assessment of acquired deficiencies, even in “normal color vision” ranges, may have useful clinical implications for functional vision testing in disease and drug/substance toxicity conditions ^22^.

This study sought to quantify the influence of age and cataractous media opacity on color and contrast vision. Color vision changes were first quantified in healthy subjects with simulated media opacity and in normal healthy patients from all age groups. A cohort of healthy patients across a wide age range was studied to evaluate color vision changes with age. Finally, color vision changes associated with cataract surgery were quantified before and after surgery.

## Methods

This study received Institutional Review Board approval from the University of California, Irvine and was conducted in accordance with the Declaration of Helsinki. Studies performed were HIPAA-compliant and all participants enrolled in the perioperative CCT testing provided informed written consent. Three separate analyses were performed to evaluate the effects of cataracts on color vision.

### Subjects and Protocol

#### Optimal CCT test conditions and test validation (neutral density filters)

CCT testing conditions were optimized to determine if mydriasis affected CCT results, and if pinhole correction could be used to account for mydriasis or presbyopia during CCT testing. Neutral density filters were evaluated to confirm reduction in CCT scores with increased filter density. Thirty-six phakic eyes of 18 subjects (10 female, 8 male) between the ages of 23-38 (25.3 ± 3.3) were recruited for this study. Testing was performed in triplicate under the following conditions: best corrected visual acuity (BCVA, no filter), distance BCVA with mydriasis (dilated), neutral density filter (ND) 03, ND 09, ND 15 (Bernell Corp), and pinholes. One eye was temporarily patched while the fellow eye underwent testing. The inclusion criteria were: (1) age ≥18 (2) no history of ocular disease or procedures (3) best corrected visual acuity of 20/20 or better in the study eye.

#### Simulated cataractous media opacity (brown filters)

Cataract-like media opacity were simulated using brown filter lenses with increasing density. The inclusion criteria were the same as described for CCT testing optimization. Six phakic eyes of 6 additional healthy subjects (5 female, 1 male) between the ages of 23-29 without any ocular history were recruited for this study. Right eyes were tested under the following conditions: no filter, filter 1, filter 2, filter 3, filter 4 (a combination of filter 2 and filter 3 in series), and “blue blocker” filter (BPI, Empire Optical). The left eye remained patched throughout testing.

#### Filter spectral transmission

The spectral transmission for the different brown filters was obtained using a compact spectrometer (CCS200/M, THORLABS, Newton, NJ). Indirect sunlight from clear skies was used to create a baseline spectrum without photobleaching the sensor. The spectra from each filter were collected while maintaining the same position and angle of the sensor. Relative transmission of each filter was calculated. The spectral emission of the CCT monitor for red, green, and blue screens were also measured.

#### Age and lens status associations with color vision

The relationship between age and color vision in phakic and pseudophakic subjects was retrospectively reviewed in healthy patients attending optometry and medical clinics. Healthy phakic (256) eyes were identified from the electronic medical record. Inclusion criteria for phakic subjects included: (1) age between 6-90 (2) no history of cataract extraction or surgery in study eye (3) no history of ocular disease in study eye other than age related cataract formation. A second analysis evaluated color vision in older patients. Forty-two phakic and 57 pseudophakic eyes were identified and evaluated with CCT. Inclusion criteria for the second older age analysis included: (1) age ≥ 50 (2) history of age-related cataract surgery in the study eye (3) no history of ocular disease aside from cataract in study eye. Testing in pseudophakic subjects was performed well after (> 3 months) the perioperative period from cataract surgery.

#### Effects of cataract on color vision

Finally, the effect of cataract was studied by evaluating CCT performance before and after cataract surgery. Thirty eyes (11 OD, 19 OS) from 25 patients (12 female, 13 male) between the ages of 51-84 were recruited for this prospective study. The inclusion criteria were: (1) age >50 (2) no history of retinal/macular pathology affecting central vision (3) completion of post-operative visit. Subjects were excluded if their visual acuity decreased by >2 lines on Snellen at their post-op month 1 visit. Visual acuity and color vision were assessed before surgery and 1-2 months after cataract extraction with intraocular lens insertion to minimize the occurrence or retinal/corneal edema that may have developed in the proximal perioperative period. At their post-op visit, pseudophakic subjects with a monofocal IOL targeted for distance vision performed CCT with near add correction, while patients with intraocular lens selection for near vision, multifocal lens or extended depth of field (EDOF) lenses wore no reading add. Information regarding IOL type and color was obtained for 30 eyes and is listed in Table 1.

**Table 1.** IOL tint and type (n=30). EDOF= extended depth of field.

| **IOL Color** | **Number** |
| --- | --- |
| Clear | 28 |
| Yellow | 2 |
| **IOL Type** |  |
| Monofocal | 23 |
| Multifocal | 4 |
| EDOF | 3 |

### Functional Testing

#### Cone Contrast Threshold

The ColorDx CCT HD (Konan Medical, Irvine, CA) is an adaptive visual function testing device that selectively stimulates retinal L-cones, M-cones, and S-cones. Using a color-calibrated anti-glare screen, a series of tumbling Landolt-C optotypes were presented in randomized directions in either decreasing or increasing steps of cone contrast against an isochromatic photopic (∼74 cd/m2) background. Subjects were asked to quickly indicate the orientation of the gap in the “C” stimulus with gated visibility of 5 seconds per stimulus as forced-choice to the subject, whether discernable or not. The optotype contrast, as LogCS, for each stimulus was recalculated based on the subject’s prior answers using a Bayesian thresholding method, the Psi-Marginal Adaptive Technique accounting for nuisance parameters like guessing and finger entry errors^20^. The next stimulus contrast value increased or decreased based upon priors (misses and correct answers) with increasingly smaller contrast steps to refine the final LogCS threshold minimizing the Standard Error. The final CCT value was derived by the Psi-marginal calculation. A summary of the individual responses as LogCS values are reported. The determined threshold also includes a unitless Rabin performance score for clinician convenience that is consistent with the tested contrasts of the original RCCT test in three ranges: >90 is normal, 75-90 indicates possible color vision deficiency (CVD), <75 indicates CVD. CCT was performed with the patients’ BCVA under photopic and monocular conditions at a distance of 2 feet.

### Statistical Analysis

Three separate analyses evaluating the effects of cataracts on color vision were performed. First, a generalized linear regression model was fit to compare the effect of each filter condition to unfiltered state for the S-cone, M-cone, and L-cone. Generalized estimating equations (GEE) were used to account for repeated measures under each condition. For the second analysis, a linear regression model fit to evaluate CCT scores for all three cone types on age and lens status was performed. As above, GEE were used to account for repeated measures since each individual received binocular testing. The third analysis explored the effects of cataract surgery on color vision using paired t-tests to assess changes in CCT scores before and after cataract extraction. All analyses were conducted using R version 1.2 ^23^.

## Results

### Optimal CCT test conditions and test validation

Greater CCT scores with least test variability was observed for the non-mydriatic state without pinholes. The greatest standard deviation for all three cones was observed in the mydriatic state with a significant reduction in M and L-cone sensitivity. Standard deviation for S-cones, M-cones, and L-cones with mydriasis were 31.6, 24.7, and 28.0 respectively. CCT scores gradually declined from baseline when a progressively more dense neutral density filter was placed between the CCT and viewer’s eye. S-cone class demonstrated the steepest decline in CCT scores with neutral density filters, while the declines in CCT scores for L and M-cones were notably less pronounced. All differences in CCT scores from baseline were statistically significant, except for the S-cone in the dilated test condition (Fig. 1).

**Figure 1.**
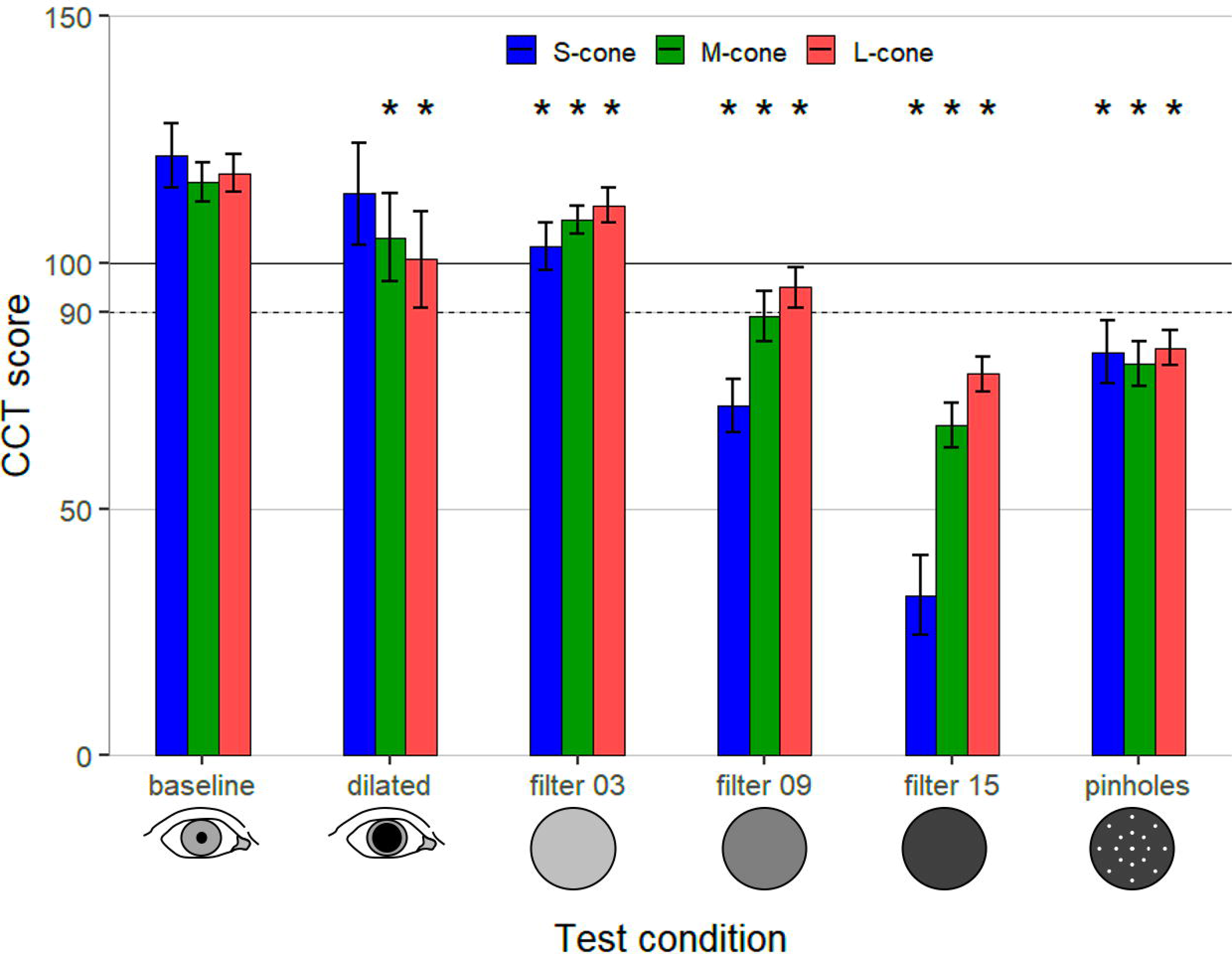
Effect of testing conditions on CCT scores, estimated via linear regression with GEE to account for repeated measurements (n=18). Error bars represent 95% confidence interval (CI) for mean CCT score and asterisks indicate statistically significant differences from baseline at p<0.05.

### Simulated cataractous media opacity (brown filters)

Six phakic eyes of 6 subjects between the ages of 23-29 (25.3 ± 2.4) without any ocular history performed CCT using brown filters simulating cataractous media opacity. Figure 2a depicts the testing system for measuring cone contrast sensitivity using different transmission filters. Figure 2b plots the measured emission of the CCT display primaries. The spectral emission ranges for red, green, and blue display primaries are: 577 nm-724 nm (peak 608 nm), 475 nm - 615 nm (peak 541 nm), and 418 nm-556 nm (peak 445 nm), respectively.

**Figure 2.**
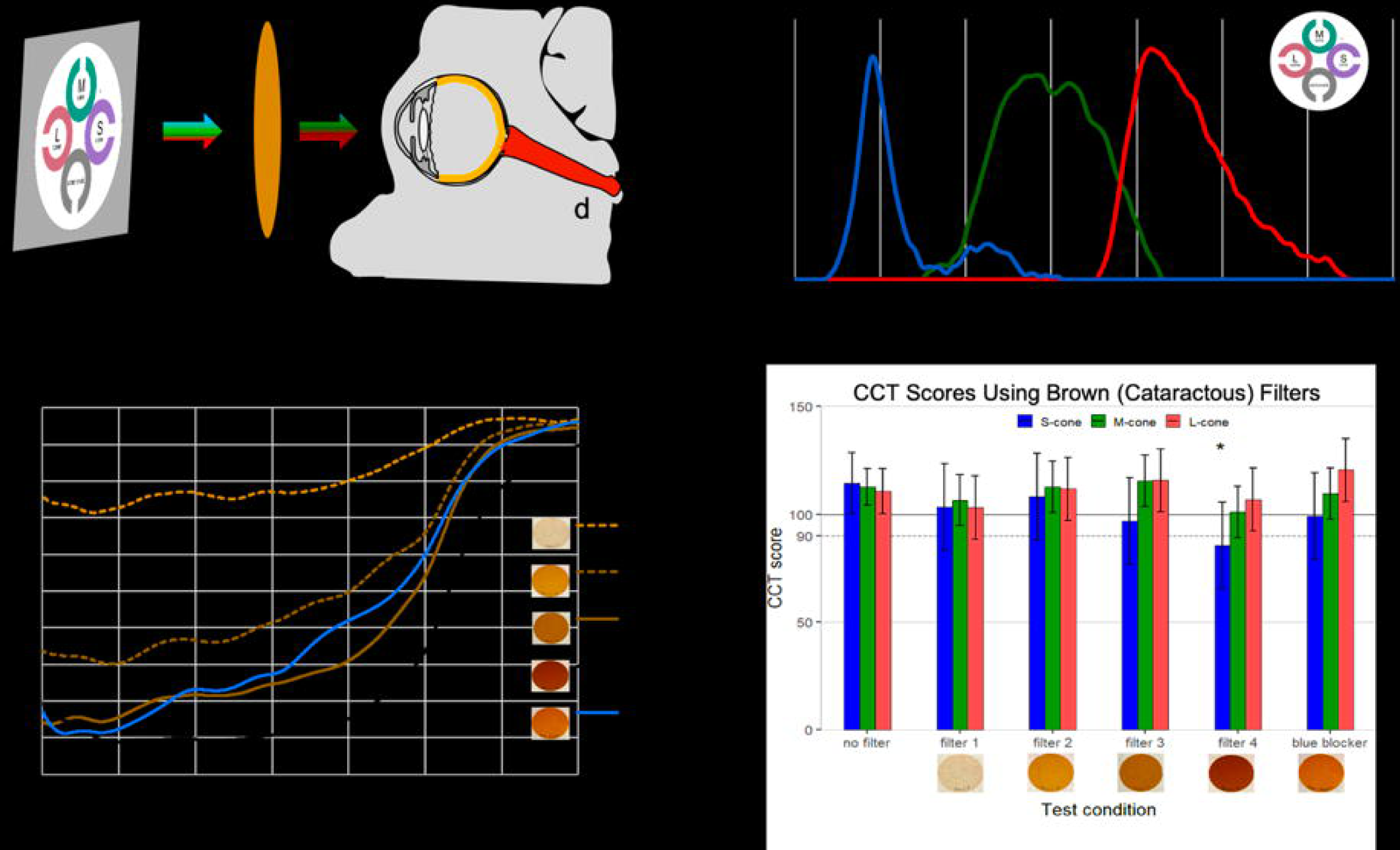
Simulating cataractous media opacity and their influence on CCT results. **(a)** Configuration for measuring cone contrast sensitivity from the ColorDx monitor, with **(b)** spectral emission of RGB light from CCT monitor, **(c)** spectral transmission of filters, and **(d)** CCT scores recorded by each subject performing the assay. Error bars represent 95% CI for mean CCT scores and asterisks indicate statistically significant differences from baseline at p<0.05 (n=6).

Figure 2c plots the transmission spectrum for each filter. Filter 1 minimally attenuates shorter wavelengths in the UV-blue spectrum and permits the greatest transmission of light in the visible spectrum of all filters tested. Filter 2, filter 3, and the blue blocker filter allow intermediate visible spectrum transmission. Filter 4 strongly attenuates the entire visible spectrum. Blue blocker transmission surpasses filter 3 transmission at wavelengths greater than 487 nm. All 5 filters permit near maximal transmission in the infrared portion of the spectrum.

Figure 2d indicates the subjective response of a test subject observing CCT through brown filters that simulate cataracts with increasing severity. When progressively dense brown filters were placed in front of the test subject’s eye, a decrease in CCT scores was seen for the S-cone class. Blue blocker produced a nominal reduction in S-cone sensitivity while the L and M-cones were nominally unaffected with larger measured confidence intervals. However, statistical significance was achieved only for the difference in S-cone sensitivity between filter 4 and no filter.

### Age-related decline in color vision

Phakic eyes (n=256) from subjects between the ages of 6-90 were included in this retrospective study based on having completed CCT performance during their clinical care. Figure 3a summarizes the effect of age on color vision in phakic eyes. CCT scores decreased for all three cone classes with increasing age. The rate of decline was greatest for the S-cones: slope (95% CI) = −1.09 (−1.23, 0.94), and least for the L-cones: slope (95% CI) = −0.66 (−0.81, −0.52).

**Figure 3.**
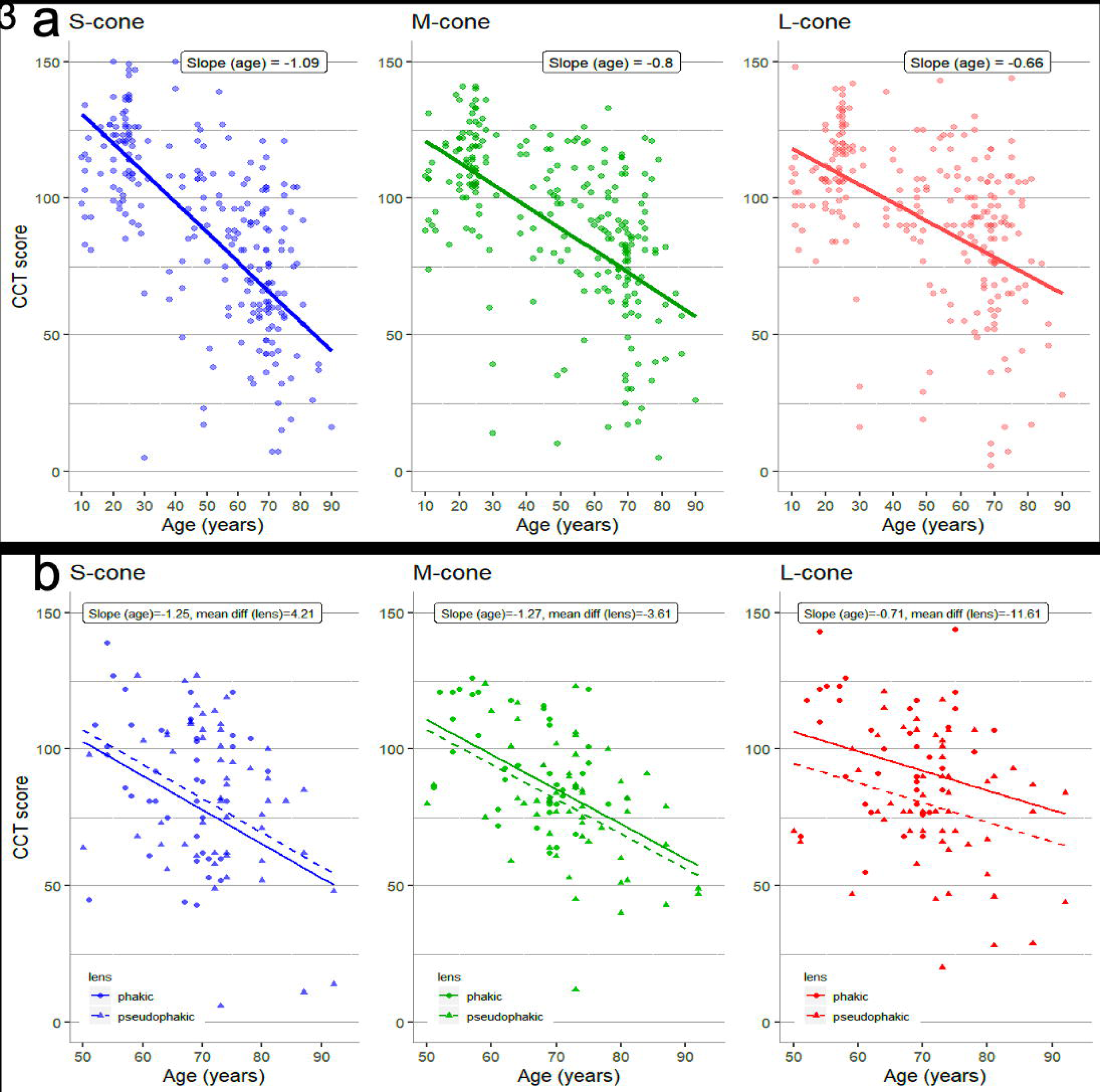
Effect of age on color vision in **(a)** phakic eyes from 6 years to 90 years old (n=256 eyes) **(b)** pseudophakic (n=57) eyes vs phakic eyes (n=42) in patients older than 50.

Figure 3b compares the effect of age on color vision between 57 pseudophakic and 42 phakic eyes in subjects 50-94 years old. CCT scores were lower in pseudophakic eyes for the M-cones and L-cones. The rate of decline was greatest for the M-cone: slope (95% CI) = −1.27 (−1.78, - 0.76), and least for the L-cones: slope (95% CI) = −0.71 (−1.37, −0.06). CCT scores were higher in pseudophakic eyes for the S-cones. The rate of decline for the S-cones was: slope (95% CI) = −1.25 (−1.98, −0.51). The differences between pseudophakic and phakic eyes for all cone classes were not statistically significant.

### Effects of cataract on color vision

Thirty eyes from 25 subjects between the ages of 51-84 (68.9 ± 6.9) were recruited to prospectively study changes in color vision with cataract surgery. Changes in CCT scores before and after cataract extraction are presented in figure 4. Mean changes between pre-op and post-op CCT scores for the S-cone, M-cone, and L-cone were 33.0 (17.8, 48.2), 24.9 (11.7, 28.1), and 22.0 (6.2, 37.8) respectively.

**Figure 4.**
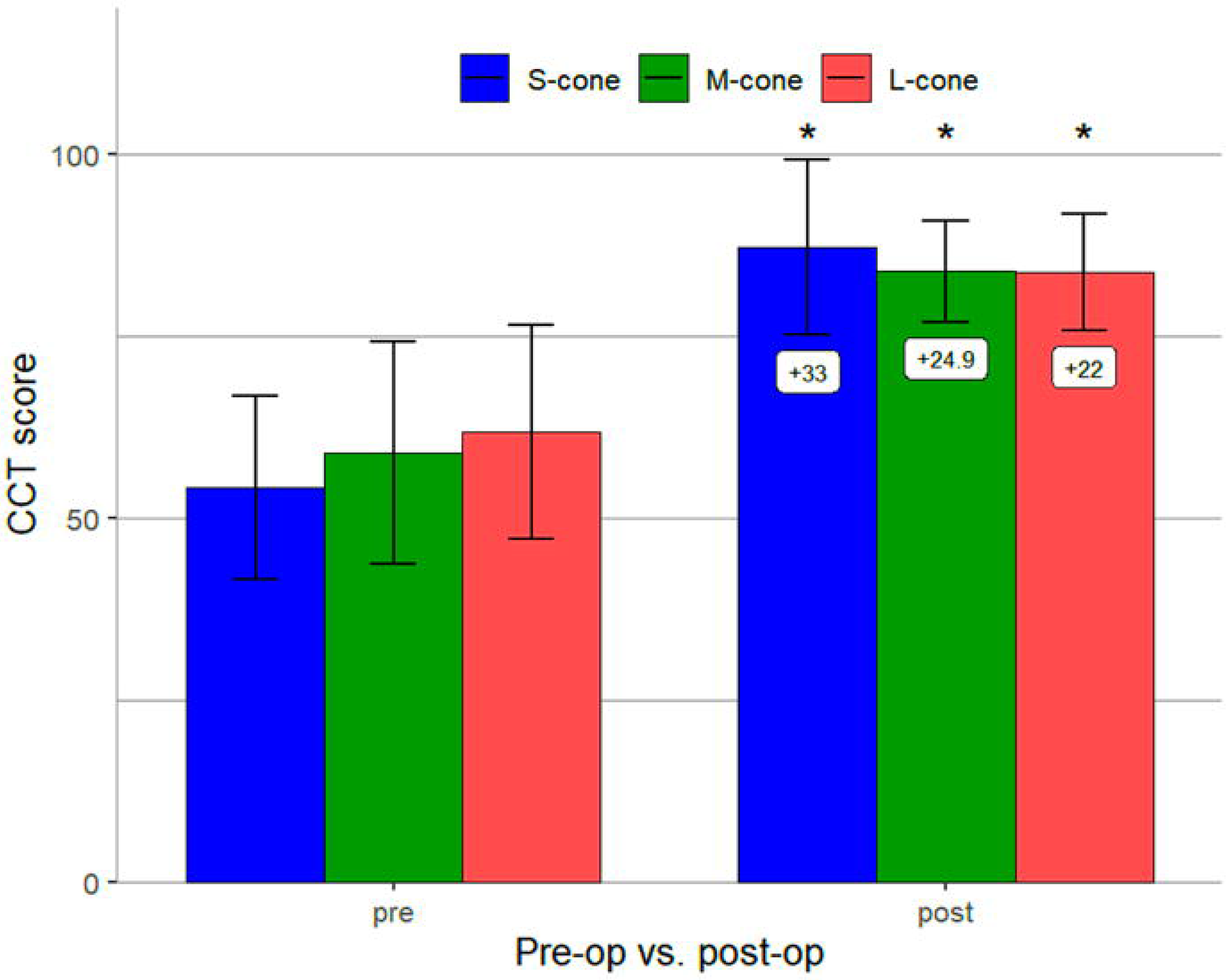
Changes in color vision before and after cataract surgery (n=25). Error bars represent 95% CI and asterisks indicate mean changes significant at p<0.05.

## Discussion

CCT is a modern quantitative color vision and contrast assay which takes approximately 7 minutes for two monocular assessments and is practical for use in clinical evaluation. Three separate analyses using CCT were performed to examine the effect of age and lens status on color and contrast vision in 3 separate scenarios: young people with simulated cataracts, populations of phakic or pseudophakic eyes seen in the course of clinical care, and subjects undergoing cataract surgery.

CCT scores significantly changed depending on testing conditions. Mydriasis produced the greatest variability in CCT results and pinhole optics reduced CCT values, indicating that CCT should be performed without mydriasis and without pinhole correction (Fig. 1). CCT scores decreased in accordance with increasing neutral filter opacity, highlighting the inverse relationship between cone contrast thresholds and lens opacity. Of note, neutral density filters (Fig. 1) produced uniform attenuation of wavelengths across the visible spectrum and should therefore not alter perceived color. However, S-cone CCT scores demonstrated greater reduction than L and M-cone classes indicating that the perceived color of an LCD display is unexpectedly changed by neutral density filters. This might be explained by the central 2 degrees of the fovea having few to no S-cones^24^. This effect was minimally reproduced using the cataract-simulating brown filters, which absorb shorter wavelengths more than longer wavelengths (Fig. 2) and was only seen with the densest brown filter. This may result from greater translucency of the darkest brown filter than the darkest neutral density filters.

CCT scores for all cone classes declined with increasing age in phakic subjects, as previously found by Fujikawa et al. S-cone exhibited the greatest rate of decline in sensitivity, highlighting the effect of lenticular aging attenuating shorter wavelengths. Alternatively, age-related decreases in color sensitivity could be attributed to increases in intraocular light scattering independent of wavelength ^21,25^, decreased cone sensitivity, misalignment of cones from photoreceptor loss, loss of nuclei in the outer nuclear layer, or other neurological factors ^2,5,15^.

Prior studies comparing color sensitivity between pseudophakic and phakic eyes found either diminished cone contrast sensitivity for the S-cones ^26^ or for all three cone classes ^2,27^ in pseudophakic subjects. However, our findings indicate greater S-cone sensitivity in pseudophakic subjects compared to healthy controls, similar to studies performed by Mantyjarvi et al^28^. This could be explained by increased filtering and decreased transmission of blue light in phakic lenses, which could outweigh the effects from photochemical damage induced by greater transmission of short-wavelength light through a clear IOL ^27^. Photochemical retinal damage would diminish color sensitivity in pseudophakic eyes, as seen for the M- and L-cones in our study. Subclinical cystoid macular edema, alternatively, could result in retinal tissue breakdown and loss of visual function^27^. However, none of the patients in our study developed pseudophakic macular edema.

In patients who underwent cataract surgery, cataracts were noted to decrease color sensitivity for all three cone classes to values below normal, with the greatest decrease seen for S-cone class. After cataract surgery, color vision increased to normal levels with the blue cone showing the greatest improvement. This finding is similar to previous studies using quantitatively discrete and continuous assays ^1,2^. Although we did not control for lens color, prior studies have shown no significant difference in color vision under photopic light conditions between blue light-filtering IOLs and UV light filtering IOLs^29^. Notably, the visible density of yellow-tinted IOLs is less remarkable than the lightest tinted brown filter studied here. To our knowledge, this is the first quantitative demonstration of this phenomenon using a standard clinical visual function test device while simulating cataractous media opacity, population-based evaluation, and perioperative assessment.

A limitation in our study is small sample size for studying progressively dense brown filter lenses. However, test subjects were healthy and reliable test takers. Another limitation is that specific time periods between filter assays were not explicitly controlled. Subjects were tested on one filter, and the subsequent filter was used 30 seconds after normal binocular vision in photopic conditions. This may have a small effect by allowing gradual dark adaptation as filters became progressively darker. However, subjects were in photopic conditions during the test and between tests with total testing time being well below 30 minutes, so dark adaptation would likely be negligible. Another limitation is that the cohort of cataract surgery patients did not exclude early disease. A history of glaucoma^2,30^, age-related macular degeneration ^31^, diabetic retinopathy^32,33^, or retinal surgeries could contribute to decreased S-cone sensitivity. To further distinguish the effect of cataract surgery on color vision for specific disease states, additional investigation could recruit disease specific sub-populations.

In conclusion, lenticular senescence and cataract formation diminish color sensitivity for all three cone classes, with the greatest decrease for S-wavelength sensitive cones. Cataract surgery can recover a significant proportion of color and contrast vision. CCT can be used to quantify the effect of cataracts and age impose on vision beyond black and white visual acuity testing. Future investigation may reveal the ways CCT testing might be used to guide recommendations for surgery or optimizing lighting conditions during activities of daily life and mitigate fall risks in the elderly ^34-36^. The degree which patients subjectively report a more ‘colorful’ and higher contrast world after cataract surgery can be quantified using CCT testing.

## Acknowledgments

The authors would like to thank Konan Medical for an unrestricted donation of ColorDx CCT HD devices.

